# Comprehensive Molecular Docking on the AlphaFold-Predicted Protein Structure Proteome: Identifying Target Protein Candidates for Puberulic Acid, a Suspected Lethal Nephrotoxin

**DOI:** 10.1101/2025.01.30.635827

**Authors:** Teppei Hayama, Rin Sugawara, Ryo Kamata, Masakazu Sekijima, Kazuki Takeda

## Abstract

This study presents a novel *in silico* screening method for identifying potential protein-ligand interactions within the complete structural proteome, as predicted by AlphaFold2, using puberulic acid as a test case. The methodology utilizes molecular docking simulations to predict interactions based on structural compatibility and binding affinity estimations. This approach highlights proteins, such as sodium/myo-inositol cotransporters, which exhibit theoretical high affinity for puberulic acid, suggesting possible competitive inhibition or mimetic interactions that could disrupt osmoregulation. It is important to clarify that these simulations provide a hypothetical framework for understanding potential interactions. The high docking scores do not directly translate to biological effects but serve as a basis for further experimental validation. This cautious interpretation acknowledges the inherent limitations of predictive modeling while maintaining confidence in the robustness of the computational process. Moreover, the exploratory nature of this study is supported by the fact that binding affinity predictions are not influenced by protein amino acid length or the predictive accuracy (pLDDT scores) provided by AlphaFold2. This independence from structural prediction metrics suggests that the docking results offer a reliable exploration of potential interactions across a wide array of proteins, making this approach a valuable asset in the fields of drug discovery and toxicological research. The ability to comprehensively identify candidate proteins with potential affinity to chemical compounds enhances the predictive capacity of toxicological assessments and supports the strategic development of safer pharmaceutical agents.

## 1. Introduction

Drug discovery is a daunting enterprise, typically spanning over a decade and costing hundreds of millions to billions of dollars, with a success rate of only 1 in 31,000 candidate compounds (Hughes et al., 2011). One major reason for this low success rate in early drug discovery phase is that most compounds manifest not only their primary effects but also unexpected side effects, particularly in off-target organs, which are notoriously difficult to predict in advance. Although animal studies are routinely used to evaluate these side effects, they primarily provide phenotypic information and make it difficult to identify the precise molecular targets underlying the observed toxicity. A notable example is the drug-related incident in Japan in 2024, involving a red yeast rice supplement suspected of causing kidney dysfunction (Ushimaru & Tominaga, 2025). Subsequent reports suggested that puberulic acid, a contaminant produced by Aspergillus during the culturing process, might be responsible. Knowledge regarding the toxicity of puberulic acid is relatively limited. Iwatsuki et al. reported its cytotoxicity against human MRC-5 cells with an IC_50_ value of 57.2 μg/ml, while Kikuchi et al. documented apoptosis-mediated cytotoxicity in human leukemia U937 cells (Iwatsuki et al., 2010; Kikuchi et al., 2024). However, its detailed mechanism of action remains unclear, underscoring the urgent need for more efficient methods to identify toxicity-related target molecules.

Historically, a variety of non-animal experimental methods, such as *in vitro* assays using cultured cells, have been employed to evaluate toxicity (Lynch et al., 2024). Although these *in vitro* assays offer higher throughput than *in vivo* animal studies, testing tens of thousands of compounds remains impractical. As an alternative, *in silico* approaches—virtual experiments performed on computers—have been explored for drug discovery. Common computational toxicity prediction techniques include quantitative structure–activity relationship (QSAR) models, which generate molecular descriptors based on chemical structure and use them to build machine learning models, and read-across methods, which infer the toxicity of an unknown compound from known toxicities of structurally similar compounds (Fisher et al., 2024). However, all of these approaches rely on structural similarities to known compounds and, therefore, struggle when confronted with novel chemical architectures or previously uncharacterized toxicity mechanisms. Although there are machine learning methods that integrate information on both ligands and their protein interaction partners, these approaches also rely heavily on existing databases of adverse effect of drugs (Raies & Bajic, 2016).

To develop new methods for predicting toxicity, it is useful to revisit how chemical substances exert toxic effects. In general, mode of action of toxicity can be described using the adverse outcome pathway (AOP) concept (Ankley et al., 2010; Vinken, 2013). The AOP framework divides toxicity into three stages: the molecular initiating event (MIE), in which the chemical binds to target biological molecules such as nucleic acids and proteins; the subsequent cellular or subcellular key events (KE); and the final adverse outcome (AO) in *in vivo*. Prediction methods for the binding of nucleic acids and chemical substances, specifically mutagenicity, are relatively well-developed. Studies have demonstrated that common machine learning techniques, such as support vector machines and gradient boosting, can predict Ames test outcomes with high accuracy (Chu et al., 2021). In contrast, proteins exhibit greater structural diversity and a wider range of functions compared to nucleic acids, making systematic understanding and prediction of their interactions and effects much more challenging and still an unmet goal, however, accurately predicting which proteins a chemical might bind to provides a viable strategy for forecasting toxicity. Molecular docking simulations—an *in silico* technique—offer a way to make such predictions (Trisciuzzi et al., 2018). These simulations place a chemical compound and a protein within a computational space and search for the most energetically favorable binding pose (Prieto-Martínez et al., 2018). Unlike *in vitro* and *in vivo* experiments, docking calculations are highly efficient, often taking only a few seconds per compound. For instance, one study screened 1,314 compounds for potential HIV protease inhibitors using the docking programs AutoDock4 and AutoDock Vina (Chang et al., 2010). However, molecular docking requires three-dimensional structures of both the chemical compounds and the proteins. Whereas three-dimensional structures of small molecules can be readily generated from their chemical formulas, experimentally determining protein structures—often comprising tens of thousands of atoms—has long required resource-intensive methods such as X-ray crystallography or cryo-electron microscopy (Bertoline et al., 2023). Recently, however, protein structure prediction techniques based on amino acid sequences, exemplified by AlphaFold2, have attained high accuracy (Jumper et al., 2021). This progress enables a comprehensive acquisition of structural proteomes, even for proteins with limited database information. In turn, the ability to perform new docking simulations on such proteins will extend toxicity predictions beyond what can be inferred from existing knowledge bases.

In this study, we implemented an *in silico* pipeline in which any chosen ligand is docked against the entire structural proteome—predicted by AlphaFold2—of an organism of interest. We then conducted an enrichment analysis on the set of proteins whose docking scores exceeded a certain threshold, thereby predicting the potential biological changes that might arise from ligand binding (Fig. 1).

**Fig. 1.**
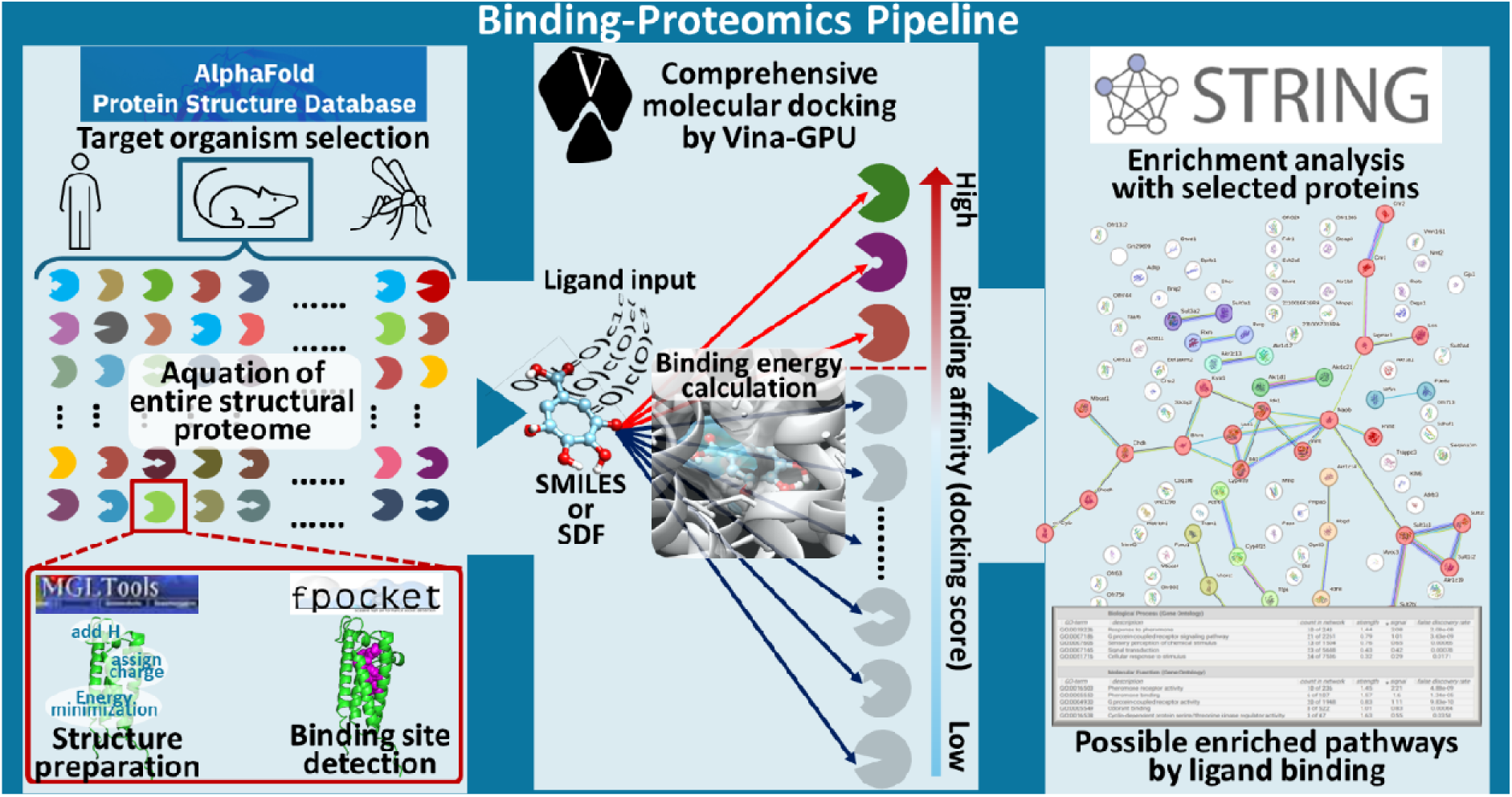
Flowchart of binding proteomics pipeline.

## 2. Materials and Methods

### 2-1. Comprehensive Acquisition and Preprocessing of Protein 3D Structure Files

A script was developed—primarily using the wget command—to download the 3D protein structure files (PDB files) for any given species from the AlphaFold Protein Structure Database (Varadi et al., 2022). As of September 16, 2024, the database provides downloadable full proteomes for 48 different species. Prior to conducting the docking simulations, all downloaded PDB files were processed with Open Babel to add polar hydrogens, assign Gasteiger charges, and convert the files into PDBQT format (O’Boyle et al., 2011). Gasteiger charges are approximate atomic charges computed from electronegativity values (Gasteiger & Marsili, 1978). PDBQT files are a modified version of the PDB format used by AutoDock Vina (Eberhardt et al., 2021; Trott & Olson, 2010). They contain the partial charges (e.g., those assigned by the Gasteiger method) as well as additional docking-related information such as hydrogen-bond donor/acceptor status and aromatic ring membership.

### 2-2. Detection of Ligand-Binding Pockets

Ligand-binding pockets for each protein were identified using Fpocket, a high-speed pocket detection program (Le Guilloux et al., 2009). Default parameters were applied. If multiple pockets were detected within a given protein, the one with the highest Pocket Score was selected as the representative ligand-binding site.

### 2-3. Molecular Docking Simulations

Docking calculations were conducted on the entire structural proteome using Vina-GPU for GPU-accelerated molecular docking (Ding et al., 2023). The binding pocket center determined by Fpocket was set as the center of the docking box. By default, the box size was set to 20 × 20 × 20 Å (adjustable as needed); in this example, a 20 × 20 × 20 Å box was employed. All other parameters for Vina-GPU were kept at their default values.

### 2-4. Enrichment Analysis

Enrichment analyses were performed using the STRING database API (Application Programming Interface) on two sets of proteins: (1) those with docking scores above a threshold (default: −8.0 kcal/mol) and (2) the top n proteins (default: n = 100) ranked by docking score (Szklarczyk et al., 2021). The version of STRING used was 11.5. First, the “Mapping identifiers” method in the STRING API was applied to convert the UniProt Accessions, associated with each PDB file, into STRING Identifiers (Bateman et al., 2023). Subsequently, the “Performing functional enrichment” method in the STRING API was used to obtain functional enrichment data. These analyses included annotations for Gene Ontology, KEGG pathways, UniProt Keywords, PubMed publications, Pfam domains, InterPro domains, and SMART domains (Consortium et al., 2023; Kanehisa et al., 2023; Letunic et al., 2021; Mistry et al., 2021; Paysan-Lafosse et al., 2023; Sayers et al., 2022).

### 2-5. Pipeline Implementation

The pipeline was designed to accept ligands not only in 3D structure file formats such as SDF or MOL2, but also as SMILES notation, which is then converted automatically. Both SDF and MOL2 are text-based file formats containing atomic coordinates and bonding information, whereas SMILES denotes molecular structures as a single line of text. When SDF or MOL2 files are provided, hydrogens are added, Gasteiger charges are assigned, and the files are converted into PDBQT format via Open Babel, following the same procedure used for protein files. In the case of SMILES input, RDKit is used to add hydrogens and generate a 3D structure via the ETKDG method before converting the file to SDF format; from there, the same procedure is applied to generate the final PDBQT files (Bento et al., 2020; Riniker & Landrum, 2015)

### 2-6. Statistical Analysis

Several statistical analyses were performed in JMP Pro 17.0 to assess the reliability of the comprehensive molecular docking approach. Specifically, the structural proteome was categorized according to Gene Ontology (GO) Slim classifications to compare docking scores across different ontologies. Pearson correlation coefficients were also computed between docking scores and two factors: pLDDT values and the protein’s amino acid sequence length.

### 2-7. Validative Flexible and Covalent Docking

Refinement of the docking results for candidate binding proteins identified in the comprehensive screening was carried out via flexible docking and covalent docking in the Schrödinger Suite. The OPLS4 force field was employed for energy minimization of both the protein and the ligand (Lu et al., 2021). Binding pockets were identified with SiteMap, and flexible docking was performed using Glide Induced Fit Docking with XP scoring and otherwise default conditions (Friesner et al., 2004). Covalent docking (CovDock) was then conducted on the poses obtained from flexible docking, designating Lys157 and Lys316 as potential covalent interaction sites (Zhu et al., 2014).

## 3. Results

### 3.1. Docking Results for Puberulic Acid Across the Full Mouse Proteome

The entire set of 21,615 mouse protein PDB files was successfully converted into PDBQT format, and Fpocket identified ligand-binding pockets in 21,581 of these structures. Docking simulations of puberulic acid against these 21,581 proteins yielded an average docking score of 5.1 (−kcal/mol), a median of 5.2, a maximum of 9.0, and a minimum of −24.8. These scores were categorized based on GO Slim terms (Figure 2). Within the “Cellular Component” GO category, the Shapiro–Wilk test indicated *p* < 0.05 for 23 out of 25 terms, which was interpreted here as evidence that the data were sufficiently close to normal distributions for most GO terms. The Levene test returned *p* = 0.0865, suggesting variance was not strictly homogeneous but was sufficiently balanced to permit an ANOVA. The ANOVA *p* value of 0.8945 indicated no statistically significant differences among GO terms in this category.

**Fig. 2.**
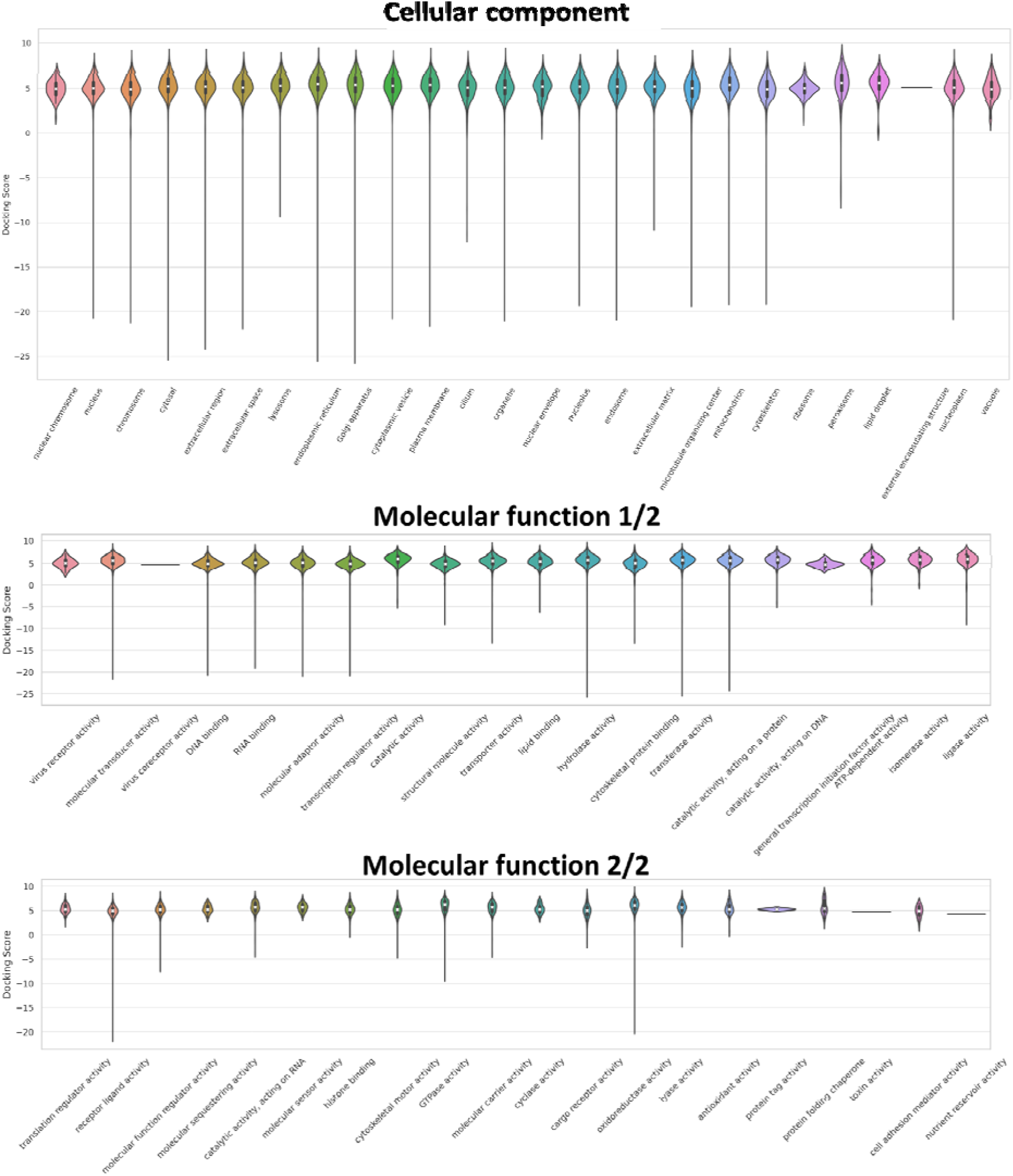
Violin plots of the docking scores for puberulic acid against the entire mouse proteome, grouped by GO Slim terms. No statistically significant difference was observed among categories in the Cellular Component ontology (p > 0.05). However, a significant difference was detected among categories in the Molecular Function ontology (p < 0.05).

In contrast, analysis of the “Molecular Function” GO category revealed that only 10 of the 36 terms had Shapiro–Wilk *p* values below 0.05, suggesting the data did not meet the assumption of normality for most terms. Furthermore, the Levene test *p* value of 0.00012805 indicated heterogeneity of variance. Consequently, the non-parametric Kruskal–Wallis test was performed, yielding *p* < 0.0001 and demonstrating a significant difference in docking scores among GO terms in the “Molecular Function” category. Figure 3A illustrates the three-dimensional surface plot of docking scores, mean pLDDT values, and amino acid sequence length for each protein. The Pearson correlation coefficient between docking score and pLDDT was 0.28, while that between docking score and amino acid length was 0.0077. Figure 3B shows a histogram of the docking scores. A total of 63 proteins (0.3% of the entire set) had docking scores of 8 or higher; these proteins were selected as candidate binding targets for further enrichment analysis.

**Fig. 3.**
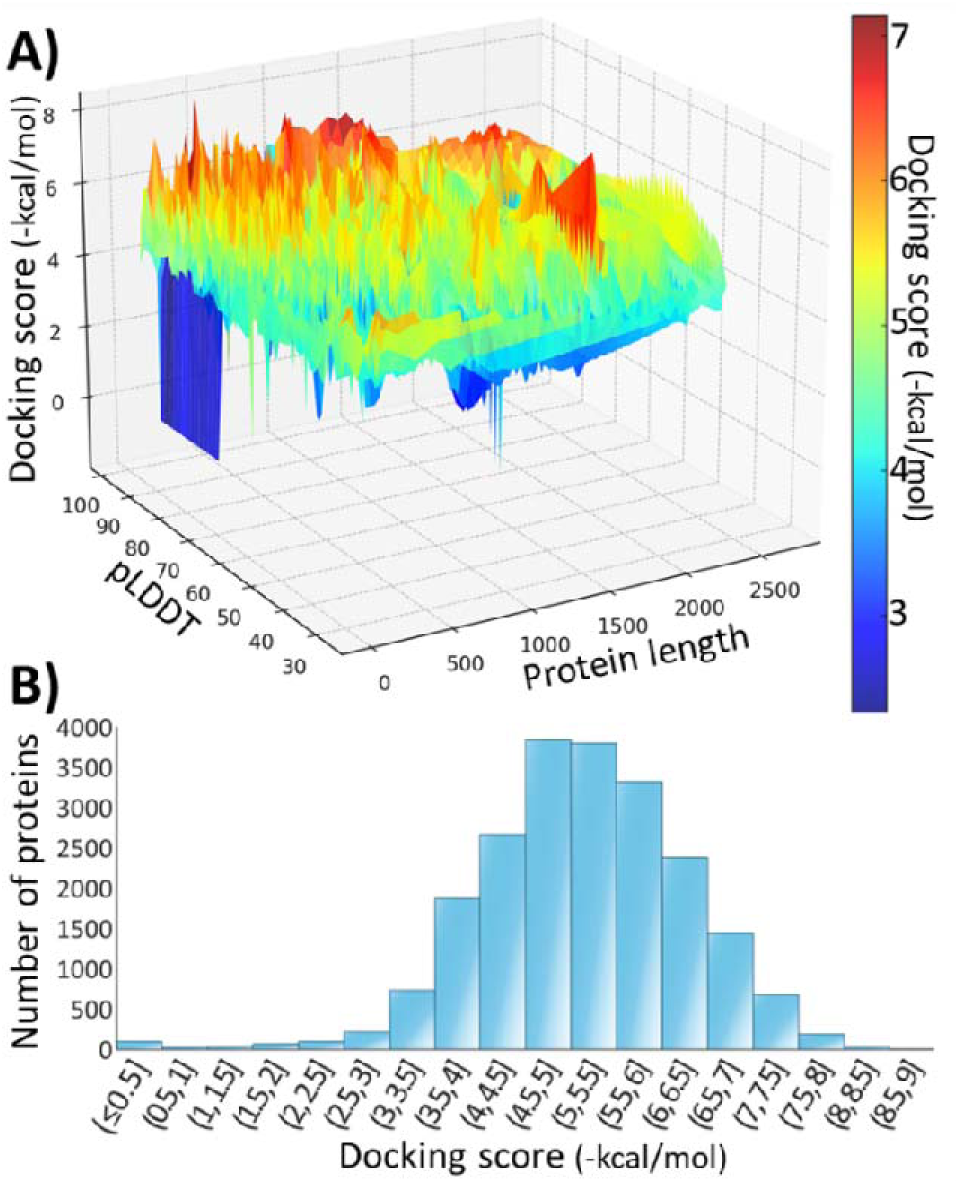
A) Three-dimensional surface plot showing docking scores for puberulic acid (versus the entire mouse proteome), pLDDT values, and amino acid sequence lengths. The pLDDT value (range: 0–100) is an AlphaFold2-derived metric that reflects protein structure prediction accuracy. Although pLDDT values are computed per amino acid, each protein is represented by its mean pLDDT value. B) Histogram of docking scores. The vertical axis represents the number of proteins in each histogram bin. A total of 63 proteins with scores ≥ 8 were subjected to subsequent enrichment analyses.

### 3.2. Enriched Pathways and Proteins

Table 1 presents the results of the enrichment analysis conducted on the high-scoring candidate binding proteins. The top-ranked pathway was “Symporter activity,” followed closely by similar terms such as “Symport” (ranked 3rd) and “Glucose:sodium symporter activity” (ranked 5th). Table 2 lists the top 20 high-scoring proteins, headed by inositol oxygenase (MIOX)—a myo-inositol–oxidizing enzyme that converts myo-inositol into D-glucuronic acid. myo-Inositol functions as a major osmolyte, transported by proteins such as sodium/myo-inositol cotransporter 2 (SMIT2, SLC5A11, rank 2 by docking score) and sodium/myo-inositol cotransporter 1 (SMIT1, SLC5A3, rank 7). Comparable analyses of the human proteome are also summarized in Tables S1 and S2. Among the proteomes of both humans and mice, only two proteins—MIOX and SMIT2—exhibited binding affinities of −8 kcal/mol or stronger for puberulic acid. In the analysis of the human proteome, MIOX ranked 4th (8.4 -kcal/mol) and SMIT2 ranked 13th (8.0-kcal/mol) among the 23,350 proteins while SMIT1 ranked 252nd (7.3 -kcal/mol).

**Table 1.** Enrichment analysis using proteins with docking scores of 8 or higher.

| Rank | ID | Pathway | FDR |
| --- | --- | --- | --- |
| 1 | GO:0015293 | Symporter activity | 0.0003 |
| 2 | GO:0036094 | Small molecule binding | 0.0003 |
| 3 | KW-0769 | Symport | 0.0005 |
| 4 | GO:0000166 | Nucleotide binding | 0.0025 |
| 5 | GO:0005412 | Glucose:sodium symporter activity | 0.0025 |
| 6 | GO:0051082 | Unfolded protein binding | 0.0025 |
| 7 | GO:0005739 | Mitochondrion | 0.0027 |
| 8 | KW:0560 | Oxidoreductase | 0.0031 |
| 9 | KW:0576 | Peroxisome | 0.0032 |
| 10 | KW:0547 | Nucleotide-binding | 0.0034 |
| 11 | GO:0003824 | Catalytic activity | 0.0039 |
| 12 | GO:0044281 | Small molecule metabolic process | 0.006 |
| 13 | MMU:425366 | Transport of bile salts and organic acids, metal ions and amine compounds | 0.0067 |
| 14 | BTO:0000671 | Kidney | 0.0068 |
| 15 | MMU:15869 | Metabolism of nucleotides | 0.0087 |
| 16 | MMU:3371568 | Attenuation phase | 0.0087 |
| 17 | MMU:6798695 | Neutrophil degranulation | 0.0087 |
| 18 | MMU:00240 | Pyrimidine metabolism | 0.0092 |
| 19 | MMU:04213 | Longevity regulating pathway - multiple species | 0.0092 |
| 20 | IPR:043129 | ATPase, nucleotide binding domain | 0.0096 |

GO: Gene ontology, KW: Uniprot keywords, MMU: Kyoto Encyclopedia of Genes and Genomes (Mus musculus), BTO: BRENDA tissue ontology, IPR: European Bioinformatics Institute's InterPro database

**Table 2.**
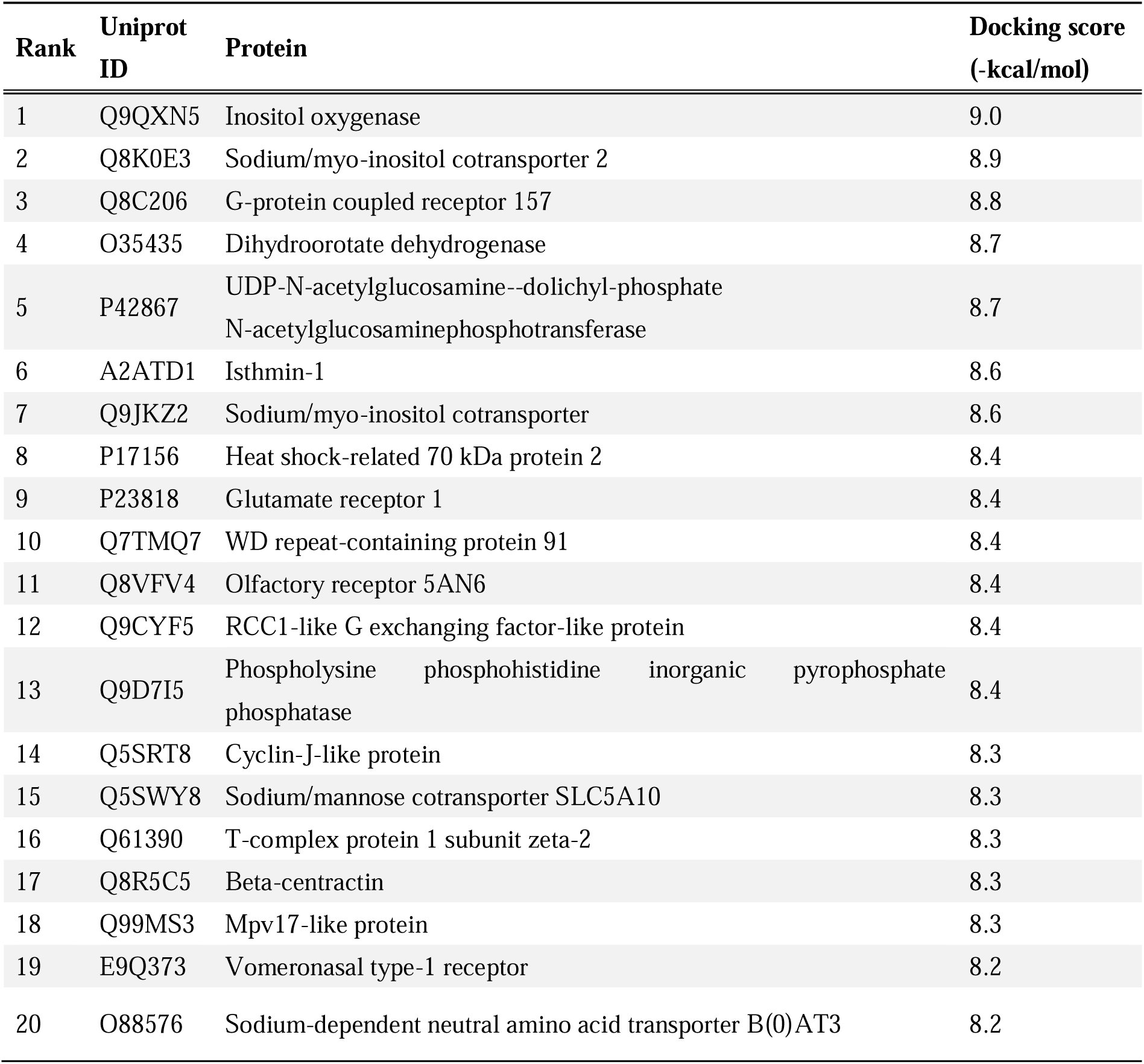
Top 20 proteins in the molecular docking score of puberulic acid to the whole mouse proteome.

### 3.3. Validative Flexible and Covalent Docking

A detailed analysis of the binding modes between puberulic acid or myo-inositol and both mouse and human MIOX, SMIT2, and SMIT1 was performed using flexible docking and covalent docking. To better reflect binding interactions in a physiological environment, flexible docking, which accounts for protein-side mobility (Glide induced fit docking), was conducted (Table 3). Puberulic acid demonstrated binding affinities comparable to those of the physiological ligand myo-inositol for all tested proteins. Observations of interacting amino acid residues revealed that lysine or arginine residues, which are capable of covalent bonding, were consistently located in close proximity to puberulic acid for all proteins (Fig. S1). Glide covalent docking was used to predict whether these residues could form imine bonds with the carbonyl group of puberulic acid via imine condensation. For all tested proteins, puberulic acid formed stable covalent bonds with the imine residues while maintaining strong binding affinities (Table 3, Fig. 4, Fig. S2, and S3).

**Fig. 4.**
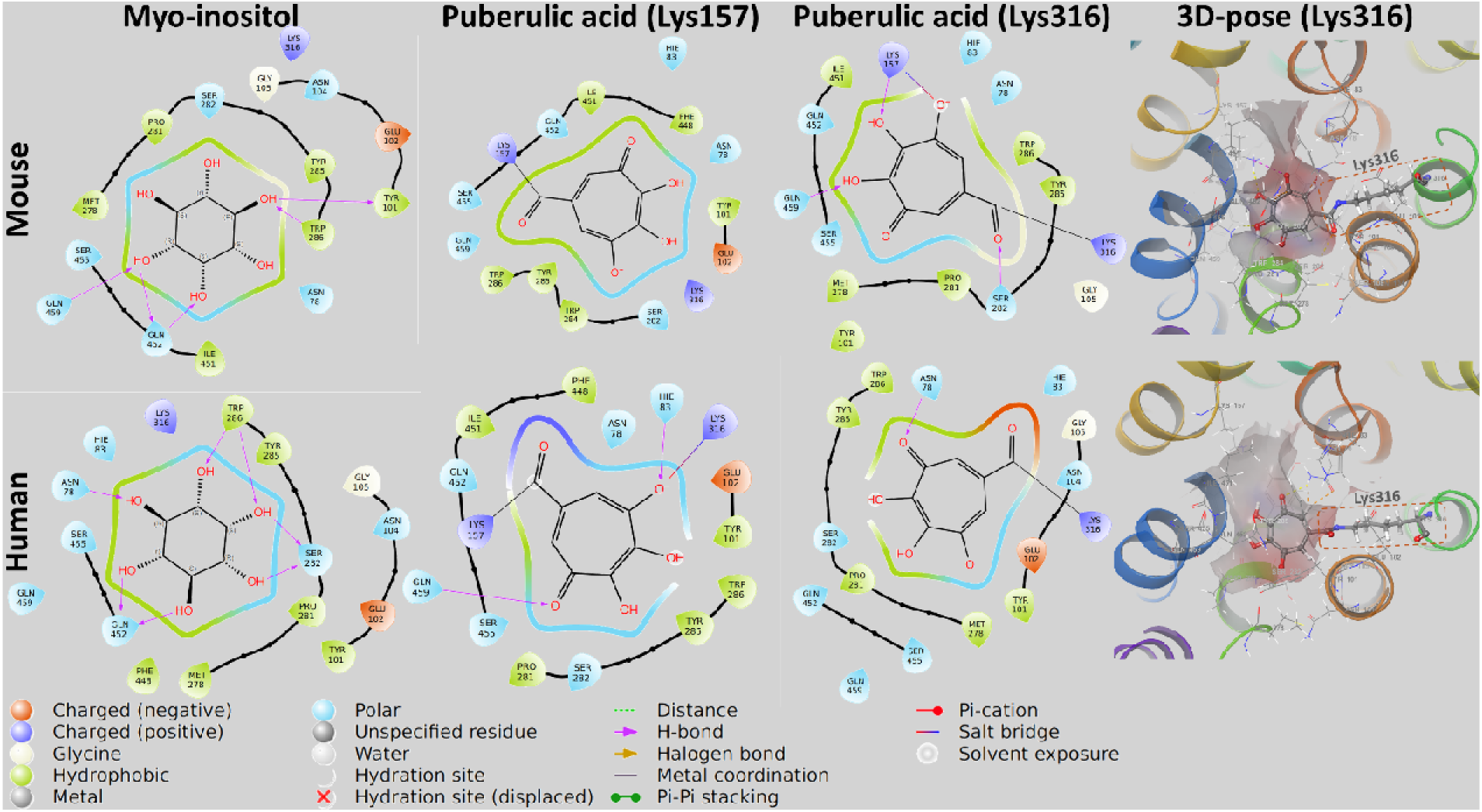
Theoretical binding-mode analysis of puberulic acid with human/mouse SMIT2 (SLC5A11), verified by flexible docking. Both the endogenous ligand myo-inositol and puberulic acid were subjected to Glide Induced Fit Docking, followed by additional covalent docking of puberulic acid with Lys157 and Lys316. The two-dimensional interaction diagrams show the key contacts between SMIT2 and each ligand, and the right panel illustrates the three-dimensional pose of puberulic acid covalently bound to Lys316.

**Table 3.** Docking scores from high-precision Glide docking (-kcal/mol).

| Ligand | Docking mode | MIOX |  | SMIT2 |  | SMIT1 |  |
| --- | --- | --- | --- | --- | --- | --- | --- |
|  |  | Mouse | Human | Mouse | Human | Mouse | Human |
| Inositol | Induced fit | 7.416 | 7.664 | 9.730 | 10.151 | 7.488 | 8.625 |
| Puberulic acid | Induced fit | 11.253 | 10.656 | 9.065 | 9.217 | 7.823 | 9.347 |
|  | Covalent—residue1 | 8.681 | 8.360 | 6.031 | 6.384 | 8.237 | 6.658 |
|  | Covalent—residue2 | 7.963 | 10.058 | 6.586 | 6.604 | 5.057 | 5.809 |

Results are shown for flexible docking (induced fit) and covalent docking for both myo-inositol and puberulic acid. Covalent docking was performed using reactive residues identified in proximity to puberulic acid during induced fit docking. For MIOX, residue1 is Arg29, and residue2 is Lys257. For SMIT1, residue1 is Lys138, and residue2 is Lys306. For SMIT2, residue1 is Lys157, and residue2 is Lys216.

## 4. Discussion

The pipeline was developed to dock a target ligand against the entire AlphaFold2-predicted proteome of any species, followed by enrichment analysis of proteins that exceed a specified docking score threshold, thereby predicting potential binding targets and their physiological effects. Because molecular docking requires the specification of a search center within the protein, the high-speed, high-accuracy Fpocket tool was employed to identify ligand-binding pockets. Among 21,615 mouse proteins, Fpocket failed to detect any ligand-binding pockets in only 34 proteins (0.16%), all of which had fewer than 100 amino acid residues (mean of 31 residues). These proteins were considered too small to form a stable binding pocket and were excluded from subsequent analyses without significantly impacting the results. Comprehensive docking simulations of puberulic acid were then carried out on the remaining 21,581 proteins. The correlation coefficient between docking scores and pLDDT (a structural prediction accuracy metric) was 0.28, indicating that the effect of AlphaFold2’s prediction accuracy on calculated binding affinities is relatively small. Furthermore, an ANOVA based on GO Slim terms in the Cellular Component category showed no significant difference, suggesting that the pipeline can comprehensively explore binding affinities regardless of subcellular localization. By contrast, an ANOVA in the Molecular Function category revealed a significant difference, implying that certain functional protein classes exhibit more specific binding. Overall, these results demonstrate the exploratory utility of the pipeline, exemplified using puberulic acid. Although docking scores are only approximate and cannot be directly converted into experimental binding constants (K_d_ or K_i_ values), an approximate calculation suggests that the threshold of −8 kcal/mol applied in this study corresponds to around 1 µM (Trott & Olson, 2010). Inspection of the puberulic acid docking score distribution revealed that only a small number of proteins reached or exceeded −8 kcal/mol, indicating that this ligand may have a high affinity for only a select subset of proteins. Although the actual number of proteins bound *in vivo* remains unclear, the observed distribution appears reasonable for screening purposes.

Among the more than 20,000 proteins in both the human and mouse proteomes, only two—MIOX and SMIT2—demonstrated strong binding affinities for puberulic acid (Table 2 and S2). This finding suggests that these proteins may work as primary toxicological targets of puberulic acid. Both MIOX and SMIT2 recognize myo-inositol as their physiological substrate. Myo-inositol has a structure in which each hydrogen atom on the carbons of a cyclohexane ring is replaced by a hydroxyl group. Puberulic acid, which contains multiple hydroxyl groups on a seven-membered ring, shares a certain degree of structural similarity with myo-inositol (Fig. 4). This structural similarity supports the possibility that puberulic acid may bind to these targets. Induced fit docking, a high-precision method that accounts for protein flexibility, is particularly suited for capturing physiologically relevant binding poses. This analysis revealed that puberulic acid binds strongly to inositol-related proteins, with SMIT2 consistently exhibiting higher binding affinities than SMIT1, aligning with trends observed in the initial screening (Table 3). Additionally, interaction analyses identified reactive lysine and arginine residues near puberulic acid within the binding pockets of MIOX, SMIT2, and SMIT1—residues that are conserved across humans and mice (Fig. S1). To evaluate the potential for covalent bond formation, covalent docking simulations were performed by integrating induced fit docking with imine condensation modeling. These simulations demonstrated that puberulic acid formed stable covalent bonds with lysine residues via imine condensation in all three binding pockets (Fig. 4, S2, and S3). Collectively, these results suggest that puberulic acid may covalently bind to inositol-related proteins, potentially disrupting their physiological functions. Both SMIT1 (SLC5A3) and SMIT2 (SLC5A11) are sodium-dependent transporters which play key roles in osmoregulation by importing myo-inositol into cells, accumulating it under high extracellular osmotic pressure to maintain osmotic balance between the intracellular and extracellular environments (Coady et al., 2002; Kwon et al., 1992) and are highly expressed in the kidney as well as in the brain and small intestine (Aouameur et al., 2007; Fu et al., 2012). In the kidney of rodents, SMIT1 is highly expressed in the renal medulla, with its expression markedly upregulated under hyperosmotic stress while SMIT2, also regulated by hyperosmotic conditions, is predominantly expressed in the brush border membrane of proximal tubules, where it plays a key role in the reabsorption of myo-inositol (Lahjouji et al., 2007; Lewis et al., 2021). Inhibition of SMIT1/2 is likely to disrupt renal osmoregulation, potentially leading to kidney injury. Interestingly, methylene myo-inositol, an inhibitor of SMIT1, is known to cause tubular degeneration—primarily in the outer medulla (Kitamura et al., 1998). This phenotype closely resembles the tubular degeneration and necrosis, as well as the Fanconi syndrome, observed in puberulic acid toxicity. MIOX is the only enzyme known to be responsible for inositol catabolism in humans and is highly expressed in the proximal tubules of the kidney (Hankes et al., 1970). In contrast to SMIT1/2, MIOX inhibition is not generally considered harmful. In fact, under hyperglycemic conditions, MIOX expression in proximal tubules is upregulated, leading to excessive inositol oxidation and increased production of reactive oxygen species (ROS). As a result, MIOX inhibition has been proposed as a potential therapeutic strategy for managing diabetic nephropathy (Tominaga et al., 2015). However, the long-term and chronic effects of MIOX inhibition on whole-body physiology, including in humans, remain poorly understood. Based on these findings, a plausible mechanism for puberulic acid–induced renal damage involves its structural similarity to myo-inositol. Puberulic acid may be taken up by SMIT1/2, forming irreversible covalent bonds with their highly reactive lysine residues, thereby inhibiting symporter activity. Loss of this critical osmoregulatory function is likely to lead to proximal tubular injury. It is important to note that all findings in this study are simulation-based, thus experimental validation for the binding affinity between the puberulic acid and SMIT1/2 and its physiological effect is required in future studies.

This pipeline can theoretically be applied to any animal species, provided that the protein sequence information is available. As previously noted, MIOX and SMIT2 were identified as the only proteins that exhibited consistently strong binding affinities for puberulic acid across both human and mouse proteomes. However, pathway analysis revealed that symporter-related pathways were significantly enriched only in mice, whereas no significant enrichment was observed in human but rather multiple terms related to GTP were listed (Table S1). This outcome stems from the high-ranking binding affinities of GTP-binding proteins such as GIMD1 and GTP-binding protein 8, along with Atlastin-1 and -2, which are GTPases involved in shaping and maintaining the tubular architecture of the endoplasmic reticulum (ER) (Table S2). These GTP-binding proteins are capable of hydrolyzing GTP in intracellular signaling pathways, thereby functioning as molecular switches. They participate in a wide range of processes, including ER and Golgi morphology, membrane trafficking, and the regulation of cell proliferation and differentiation signals. Additionally, many members of the GTP-binding protein family are deeply involved in immune regulation (Hall, 1990; Kaziro et al., 1991). Puberulic acid may mimic GTP in such a way that it modulates inflammation and immune responses through GTP-dependent signaling. These findings underscore the importance of examining multiple species. The AlphaFold database already provides ready-made ZIP files containing cross-species structural proteomes for 48 representative model organisms, and this pipeline also makes use of those files. However, by modifying the script to download structures sequentially based on species name searches, the pipeline can be extended, in principle, to virtually any species.

As a limitation of the present study, the analysis is restricted to an energy-based molecular docking approach followed by enrichment analysis and does not constitute a direct prediction of side effects. Rather, the pipeline is intended for use as a component of binding proteomics in toxicological research, complementing transcriptomic or proteomic analyses by providing candidate binding proteins and pathways to toxicologists. A related approach involves using large-scale molecular docking data on compound–protein pairs to build training sets for side-effect prediction (Sawada et al., 2024). However, such methods are often influenced by existing knowledge of compound toxicity, making them less suitable for identifying novel target proteins. By contrast, the pipeline described here provides purely energy-based rankings that may facilitate new discoveries. The fact that SMIT1 and SMIT2 have not been well recognized as a nephrotoxicity target supports the pipeline’s potential utility as an energy calculation-based *in silico* omics analysis. Various methods exist for molecular docking and ligand-binding pocket detection, each with its own advantages and limitations. It is important to acknowledge that no current approach can perfectly predict binding poses with absolute accuracy. This pipeline employs Vina-GPU and Fpocket to achieve high-speed, comprehensive molecular docking calculations, further benchmarking using well-characterized compounds with known targets is needed. Additionally, the pipeline adopts only the highest-ranked pocket identified by Fpocket in order to control computational complexity, despite the possibility that some ligands may bind to allosteric sites. Future work may involve increasing the accuracy of the comprehensive docking calculations and refining methods for inferring biological changes from the results. Techniques such as RoseTTAFold-all-atom and AlphaFold3, which predict complexes involving ligands and nucleic acids, have already been developed; evaluating their applicability to toxicity prediction may also be worthwhile—although current licensing terms preclude docking calculations using AlphaFold3 structures (Abramson et al., 2024; Krishna et al., 2024). Since the advent of AlphaFold2, structural biology has undergone rapid and transformative advancements, not only in its core methodologies but also in its widespread application across various fields of biological research. The approach proposed in this study, termed “Binding Proteomics,” is expected to further accelerate the integration of structural biology into toxicology, enhancing our ability to predict and understand toxicity at the molecular level.

## Conclusion

This study proposes an *in silico* screening method for identifying candidate binding target proteins via comprehensive molecular docking calculations of an organism’s structural proteome. An analysis focusing on puberulic acid—a compound that has raised significant public health concerns in Japan due to its potential for lethal kidney damage—highlighted the sodium/myo-inositol cotransporter as a prominent target candidate. These findings underscore the need for further experimental investigations to validate the toxicological effects of puberulic acid.

## Data availability

The source code implementing this pipeline is available at https://github.com/toxtoxcat/reAlldock.

## Acknowledgments

This study was supported by Grants[in[Aid for Scientific Research from the Ministry of Education, Culture, Sports, Science and Technology of Japan awarded to K. Takeda (23K14091). This research was supported by Platform Project for Supporting Drug Discovery and Life Science Research (Basis for Supporting Innovative Drug Discovery and Life Science Research (BINDS)) from AMED under Grant Number JP23ama121027 awarded to M. Sekijima.

## Supplementary materials

Table S1. Enrichment analysis using proteins with docking scores of 8 or higher

Table S2. Top 20 proteins in the molecular docking score of puberulic acid to the whole human proteome

Fig. S1. Interaction diagrams of induced-fit docking for MIOX, SMIT2, and SMIT1 in human and mouse.

Fig. S2. Interaction diagrams of covalent docking for SMIT1 in human and mouse.

Fig. S3. Interaction diagrams of covalent docking for MIOX in human and mouse.

