## Supplementary materials for "Comprehensive Molecular Docking on the AlphaFold-Predicted Protein Structure Proteome: Identifying Target Protein Candidates for Puberulic Acid, a Suspected Lethal Nephrotoxin"

Table S1. Enrichment analysis using proteins with docking scores of 8 or higher

Table S2. Top 20 proteins in the molecular docking score of puberulic acid to the whole human proteome

Fig. S1. Interaction diagrams of induced-fit docking for MIOX, SMIT2, and SMIT1 in human and mouse.

Fig. S2. Interaction diagrams of covalent docking for SMIT1 in human and mouse.

Fig. S3. Interaction diagrams of covalent docking for MIOX in human and mouse.

Table S1. Enrichment analysis using proteins with docking scores of 8 or higher

| **Rank** | **ID** | Pathway | FDR |
|---|---|---|---|
| 1 | KW-0342 | GTP-binding | 0.0056 |
| 2 | IPR006073 | GTP binding domain | 0.0056 |
| 3 | IPR027417 | P-loop containing nucleoside triphosphate hydrolase | 0.0146 |
| 4 | KW-0547 | Nucleotide-binding | 0.0169 |
| 5 | GO:0098826 | Endoplasmic reticulum tubular network membrane | 0.0252 |
| 6 | GO:0000166 | Nucleotide binding | 0.0332 |
| 7 | GO:0005041 | Low-density lipoprotein particle receptor activity | 0.0332 |
| 8 | GO:0005525 | GTP binding | 0.0332 |
| 9 | GO:0016787 | Hydrolase activity | 0.0332 |
| 10 | GO:0017111 | Nucleoside-triphosphatase activity | 0.0332 |
| 11 | GO:0032555 | Purine ribonucleotide binding | 0.0332 |
| 12 | GO:0034185 | Apolipoprotein binding | 0.0332 |
| 13 | GO:0035639 | Purine ribonucleoside triphosphate binding | 0.0332 |
| 14 | GO:0036094 | Small molecule binding | 0.0332 |
| 15 | GO:0043167 | Ion binding | 0.0332 |
| 16 | GO:0043168 | Anion binding | 0.0332 |
| 17 | GO:0097367 | Carbohydrate derivative binding | 0.0332 |
| 18 | GOCC:0098826 | Endoplasmic reticulum tubular network membrane | 0.0343 |

Table S2. Top 20 proteins in the molecular docking score of puberulic acid to the whole human proteome.

| **Rank** | **Uniprot ID** | **Protein** | **Docking score (-kcal/mol)** |
|---|---|---|---|
| 1 | Q07954 | Prolow-density lipoprotein receptor-related protein 1 (LRP-1) | 8.6 |
| 2 | O75581 | Low-density lipoprotein receptor-related protein 6 (LRP-6) | 8.4 |
| 3 | P0DJR0 | GTPase IMAP family member GIMD1 | 8.4 |
| 4 | Q9UGB7 | Inositol oxygenase | 8.4 |
| 5 | P36915 | GTP-binding protein HSR1 | 8.2 |
| 6 | Q86YJ6 | Threonine synthase-like 2 (TSH2) | 8.2 |
| 7 | O00148 | ATP-dependent RNA helicase DDX39A | 8.1 |
| 8 | Q12873 | Chromodomain-helicase-DNA-binding protein 3 (CHD-3) | 8.1 |
| 9 | Q8N3Z3 | GTP-binding protein 8 | 8.1 |
| 10 | Q8NHH9 | Atlastin-2 | 8.1 |
| 11 | Q96GD0 | Chronophin (Pyridoxal phosphate phosphatase) | 8.1 |
| 12 | Q9NVN8 | Guanine nucleotide-binding protein-like 3-like protein | 8.1 |
| 13 | Q8WWX8 | Sodium/myo-inositol cotransporter 2 | 8.0 |
| 14 | P11021 | Endoplasmic reticulum chaperone BiP (HSP70 family protein 5) | 8.0 |
| 15 | P04040 | Catalase | 8.0 |
| 16 | Q8WXF7 | Atlastin-1 (GTP-binding protein 3) | 8.0 |
| 17 | Q9NR77 | Peroxisomal membrane protein 2 | 8.0 |
| 18 | Q9NRR2 | Tryptase gamma (Serine protease 31) | 8.0 |
| 19 | O43488 | Aflatoxin B1 aldehyde reductase member 2 (Succinic semialdehyde reductase) | 7.9 |
| 20 | P04075 | Fructose-bisphosphate aldolase A | 7.9 |

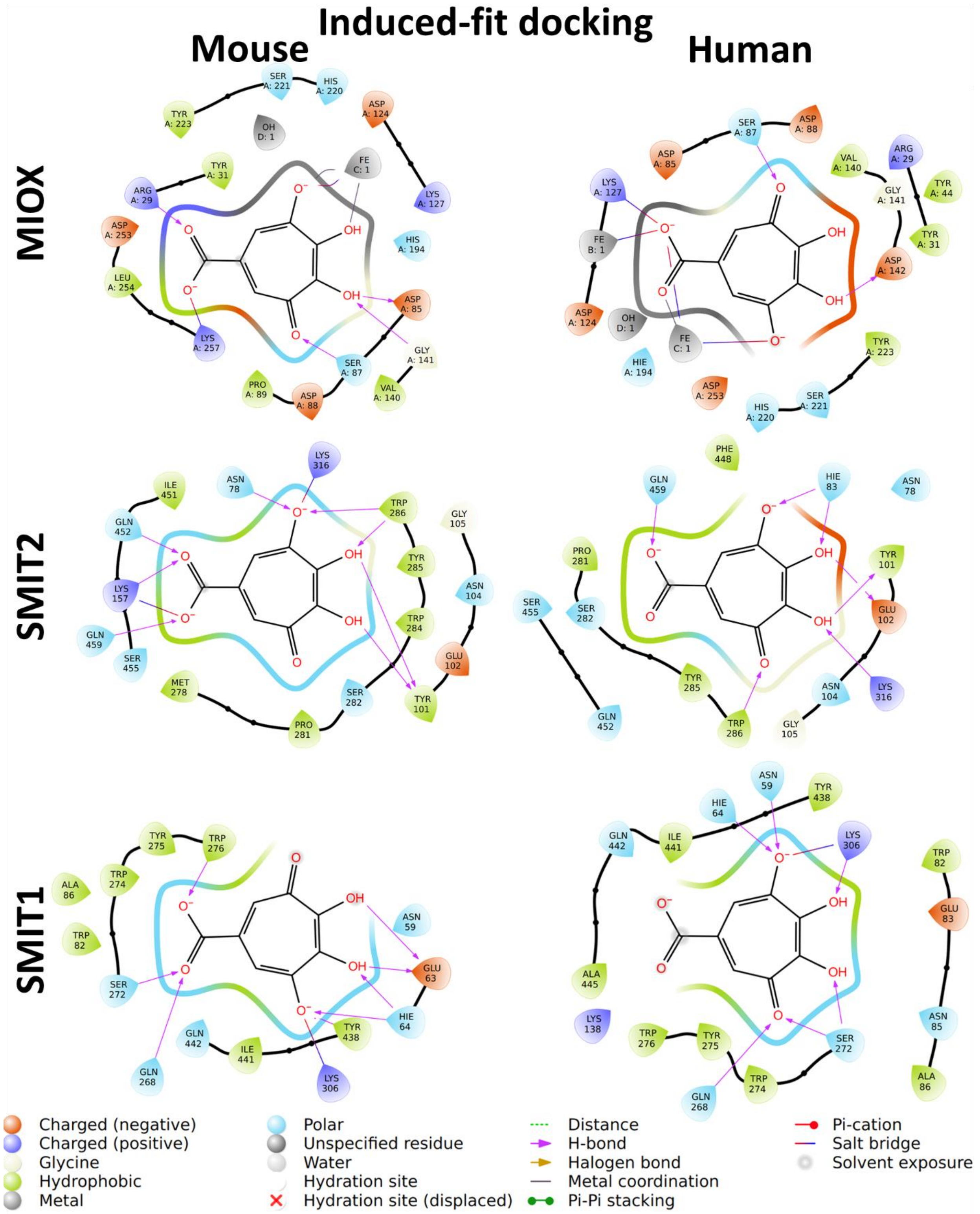


Fig. S1. Interaction diagrams of induced-fit docking for MIOX, SMIT2, and SMIT1 in human and mouse.

28 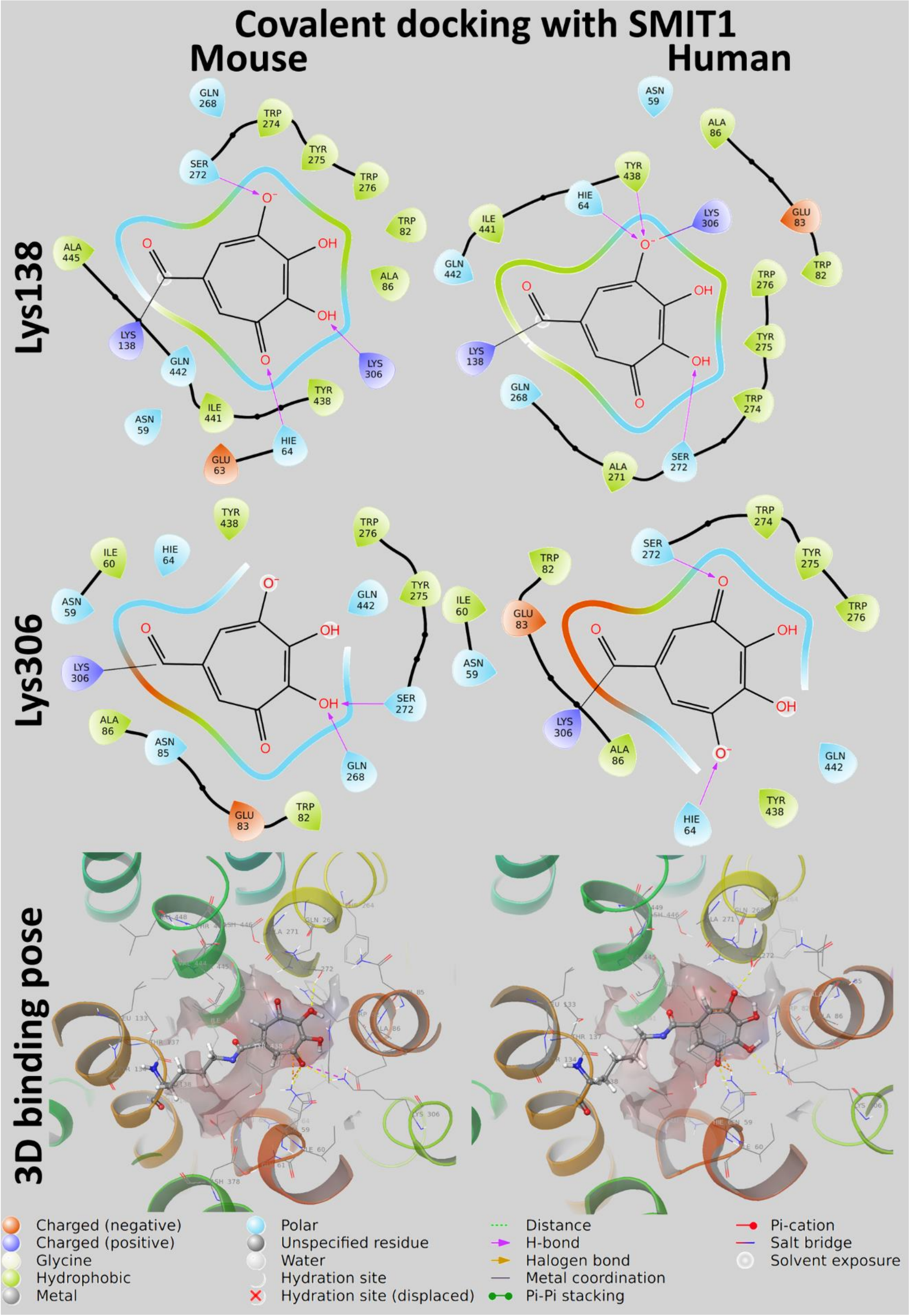


29 Fig. S2. Interaction diagrams of covalent docking for SMIT1 in human and mouse.

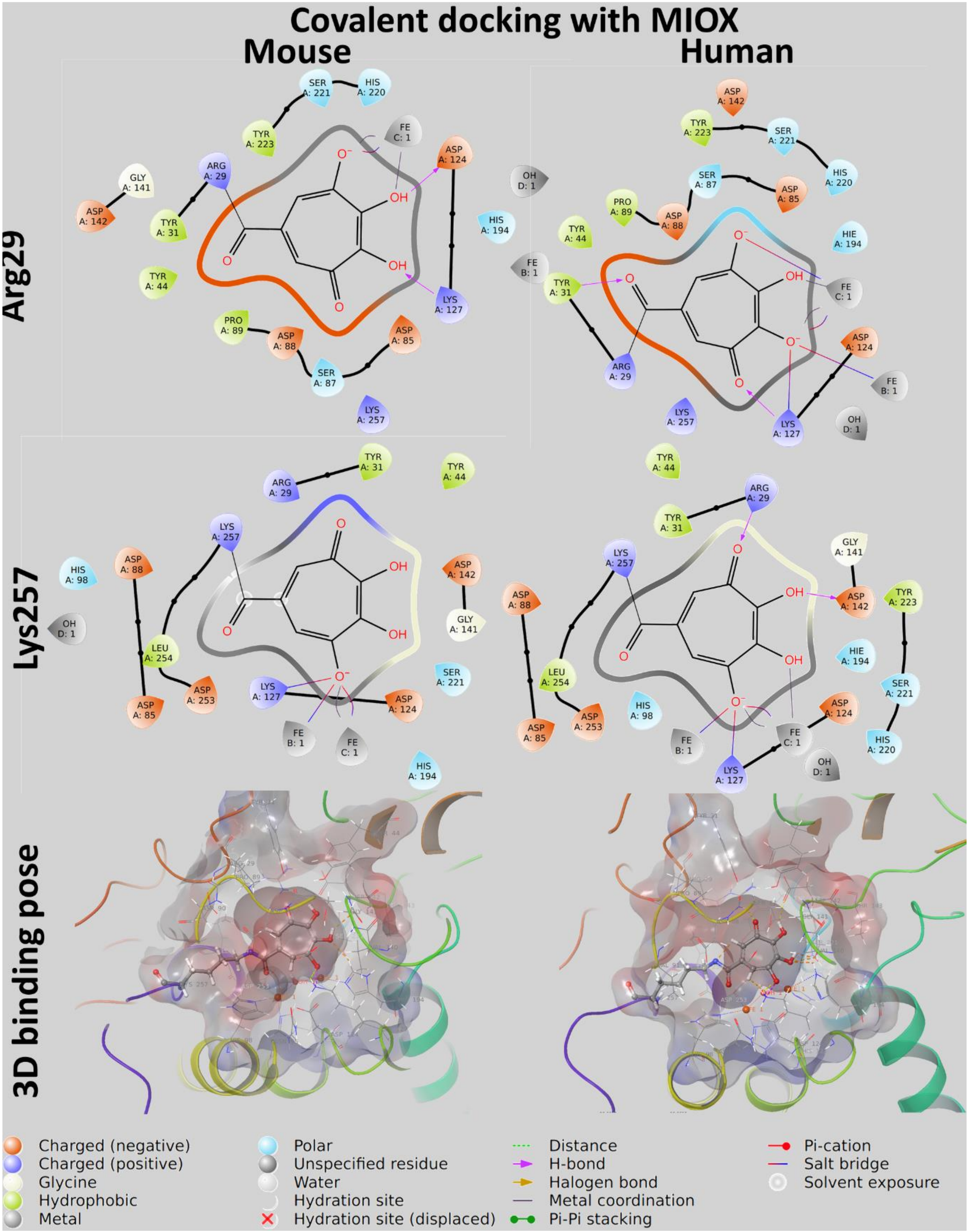


Fig. S3. Interaction diagrams of covalent docking for MIOX in human and mouse.
